# Comparison of HSV-1 strains circulating in Finland demonstrates the uncoupling of whole-genome relatedness and phenotypic outcomes of viral infection

**DOI:** 10.1101/424408

**Authors:** Christopher D. Bowen, Henrik Paavilainen, Daniel W. Renner, Jussi Palomäki, Jenni Lehtinen, Tytti Vuorinen, Peter Norberg, Veijo Hukkanen, Moriah L. Szpara

**Author notes:** equal contribution of DWR and HP. equal contribution of MLS and VH. Corresponding Author: Moriah L. Szpara, Dept. of Biochemistry & Molecular Biology, The Huck Institutes of the Life Sciences, W-208 Millennium Science Complex (MSC), Pennsylvania State University, University Park, PA 16802 USA. Author emails: Christopher D. Bowen, Henrik Paavilainen, Daniel W. Renner, Jussi Palomäki, Jenni Lehtinen, Tytti Vuorinen, Peter Norberg, Veijo Hukkanen, Moriah L. Szpara.

## Abstract

A majority of adults in Finland are seropositive carriers of herpes simplex viruses (HSV). Infection occurs at epithelial or mucosal surfaces, after which virions enter innervating nerve endings, eventually establishing lifelong infection in neurons of the sensory or autonomic nervous system. Recent data have highlighted the genetic diversity of HSV-1 strains, and demonstrated apparent geographic patterns in strain similarity. Though multiple HSV-1 genomes have been sequenced from Europe to date, there is a lack of sequenced genomes from Nordic countries. Finland’s history includes at least two major waves of human migration, suggesting the potential for diverse viruses to persist in the population. Here we used HSV-1 clinical isolates from Finland to test the relationship between viral phylogeny, genetic variation, and phenotypic characteristics. We found that Finnish HSV-1 isolates separated into two distinct phylogenetic groups, potentially reflecting historical waves of human (and viral) migration into Finland. Each HSV-1 isolate harbored a distinct set of phenotypes in cell culture, including differences in virus production, extracellular virus release, and cell-type-specific fitness. Importantly, the phylogenetic clusters were not predictive of any detectable pattern in phenotypic differences, demonstrating that whole-genome relatedness is not a proxy for overall viral phenotype. Instead, we highlight specific gene-level differences that may contribute to observed phenotypic differences, and we note that strains from different phylogenetic groups contain the same genetic variations.

**Importance:** Herpes simplex viruses (HSV) infect a majority of adults. Recent data have highlighted the genetic diversity of HSV-1 strains, and demonstrated apparent genomic relatedness between strains from the same geographic region. We use HSV-1 clinical isolates from Finland to test the relationship between viral genomic and geographic relationships, differences in specific genes, and characteristics of viral infection. We found that viral isolates from Finland separated into two distinct groups of genomic and geographic relatedness, potentially reflecting historical patterns of human and viral migration into Finland. These Finland HSV-1 isolates had distinct infection characteristics in multiple cell types tested, which were specific to each isolate and did not group according to genomic and geographic relatedness. This demonstrates that HSV-1 strain differences in specific characteristics of infection are set by a combination of host cell type and specific viral gene-level differences.

## Introduction

Human herpes simplex virus 1 (HSV-1; Family: *Herpesviridae*, Genus: *Simplexvirus*, species: *Human herpesvirus 1*) has been recognized as a cause of major human disease since the era of Hippocrates (1), and to this day there is no effective vaccine (2). Approximately 50% of adults in Finland are seropositive for HSV-1 (3–5), and 13% are seropositive for HSV-2 (3, 6). HSV infects via epithelial or mucosal surfaces, after which it is taken up by nerve endings and establishes a lifelong infection in neurons of the sensory or sympathetic nervous system. From these neuronal sites the virus can reactivate, transit via nerve endings back to the skin surface, and re-initiate skin or mucosal shedding at the same site as the original infection. In addition to recurrent epithelial lesions, HSV also causes infectious keratitis, worsens acquisition and shedding rates for human immunodeficiency virus (HIV), and can progress to cause rare but life-threatening encephalitis (1).

Recent comparative genomics studies of HSV-1, from our lab and others, have demonstrated that HSV-1 strains from unrelated individuals can differ in 2-4% of the viral genome (7–13). HSV-1 has a large DNA genome of ∼152 kilobases, encoding >75 proteins (14, 15). Prior comparisons of HSV-1 strains based on single-gene or whole-genome sequencing have found at least three major clades that cluster geographically, with evidence of recombination between these groups (16–18, 9). However these studies have focused almost exclusively on the quantification of sequence diversity, without connecting these observations to experimental measures of viral fitness and/or virulence. Several recent examples have demonstrated the usefulness of comparing closely related virus strains to explore the effects of minor sequence differences on phenotypic outcomes (19–22, 10). To date, publications examining the genome-wide diversity of HSV-1 in diverse clinical isolates have not connected these observations to any classic measures of viral fitness, such as viral replication in cells, or antiviral drug resistance (23–25). Based on the decades of prior research leading to all current antiviral approaches for HSV-1, we anticipate that a connection between viral genetic data, cell-based measures of viral fitness, and in vivo measures of viral pathogenesis will be crucial to drive the next generation of viral therapeutics. Examining the phenotypic differences displayed by HSV-1 strains in culture provides an opportunity to explore the scope of these differences and test whether or not they are linked to previously observed patterns of geographic diversity.

Finland has a unique history that has led to a relatively homogeneous and stable population (26–28), providing a unique view on the evolution of viruses that are persistent in the population. There has been gene flow contributing to human population of Finland: both from the east, and from the south (southwest) (26–28). The Eastern migrations from Siberia began over 3500 years ago, and allele sharing with modern East Asian populations can be observed even in present day Finns (27, 28). Finland also has one of the best genealogical databases in the world, which in combination with computerized medical records and a high rate of patient participation, has led to many recent advances in human medical genetics (29, 30). This may enable future studies to explore the co-evolution of human and viral genetic variation in this population.

We have characterized ten HSV-1 strains isolated from a random subset of Finland clinic visits, and compared their growth properties, drug resistance, and other phenotypic features. Full genome sequences of these viruses were compared to each other and to other previously described strains of HSV-1, to reveal patterns of inter-host and intra-host variability. While two phylogenetic sub-groups were detected at a genomic level, these sub-groups were not predictive of any detectable pattern in the observed phenotypic differences, demonstrating that whole-genome relatedness is not a proxy for viral phenotype. We anticipate that these data will aid in efforts to develop improved sequence-based antiviral therapies, by providing additional data on conserved vs. divergent areas of the HSV-1 genome, and contribute to development of a vaccine against HSV infections based on attenuation of these or related clinical isolates (31). These data present an opportunity to explore the diversity of chronic herpesviruses in the Finland population, and to lay the foundation for future studies that explore the connections between viral genetic differences, host genetic predispositions, and their potential relationship(s) to clinical outcomes of HSV-1 infection.

## Methods

### Virus isolates and virus stock propagation

Clinical isolates were obtained from anonymous coded diagnostic samples from herpes lesions representing currently circulating viral strains in Finland (**Table 1**). The virus diagnostic unit of the Turku University Hospital receives viral culture samples from the whole country except for the Helsinki-Uusimaa region (capital Helsinki area and the South-Eastern Finland). Recent immigrations to Finland were not represented in the sample material. Approval for the study of anonymous HSV isolates was provided by the Turku University Central Hospital (permit # J10/17).

**Table 1:** The Finnish HSV-1 strains used for genome sequence comparisons.

| Sample | Gender | Code | Age | Lesion or diagnosis | Acyclovir sensitivity, IC50 value (µg/ml) |
| --- | --- | --- | --- | --- | --- |
| H1211 | female | F-11 | 21 | blister | 0.25 |
| H1215 | male | M-15 | 5 | blister | 0.03 |
| H12113 | female | F-13 | 29 | blister | 0.11 |
| H12114 | female | F-14g | 23 | genital blister | 0.50 |
| H12117 | female | F-17 | 57 | blister | 0.15 |
| H12118 | female | F-18g | 19 | genital blister | 0.13 |
| H1311 | female | F-11l | 59 | blister (lip) | 0.31 |
| H1312 | male | M-12 | 39 | blister | 0.13 |
| H1412 | female | F-12g | 28 | genital lesion | 0.09 |
| H15119 | male | M-19 | 36 | blister | 0.15 |

\*The IC50 value for HSV-1 delta305 (a TK-negative, ACV-resistant control) was 8.08 and for the reference strain HSV-1(17+) it was 0.13. The limit of resistance was $\geq 2.0$ µg/ml.

An immunoperoxidase-rapid culture assay (32) was used to type the viruses as HSV-1, which was confirmed by a HSV type-specific gD (US6) gene-based PCR test (33). Viruses were initially propagated on Vero cells (African green monkey kidney cells; ATCC), maintained in Dulbecco’s MEM with 2% fetal calf serum and gentamycin. A stock was made by addition of 3 ml of 9% autoclaved skimmed milk (Valio, Finland) on to 5 ml of the culture medium and subsequent freezing. The cells and the medium were collected and combined upon thawing, and were frozen and thawed for two further rounds before aliquoting. The viral titer was determined by plaque titration on Vero cells as described before (34). Parallel aliquots were used for further viral culture studies and for preparation of viral nucleocapsid DNA.

### Statistical methods

For viral production data, SPSS Statistics 20 (IBM, Armonk NY, USA) software was use to perform statistical analyses. Non-parametric Mann-Whitney U-test was used to calculate statistical significances. The threshold for significance was set to p<0.05.

### Viral genomic DNA isolation

The viral genomic DNA was prepared from isolated viral nucleocapsids as described previously (7, 35). In brief, viral stock collected from the first or second passage in Vero cells was used to infect ∼1×10^8^ HaCaT cells (Department of Dentistry, University of Turku (36)) at a multiplicity of infection (m.o.i.) of 0.1-1 plaque-forming units (PFU)/cell, and the infection was allowed to proceed to completion at +35 °C (1-3 days). The cells were collected, washed with PBS and suspended to LCM buffer (0.125 M KCl, 30 mM Tris pH 7.4, 5 mM MgCl_2_, 0.5 mM EDTA, 0.5% Nonidet P-40; with 0.6 mM beta-mercaptoethanol). After two successive extractions with Freon (1,1,2-trichloro-1,2,2-trifluoroethane, Sigma-Aldrich), the extracts were added on top of layers in LCM buffer with 45% and 5% glycerol and ultracentrifuged at 77 000 x g for 1 h at +4°C (Beckman Coulter SW41Ti rotor). Viral nucleocapsids were recovered from the bottom of the ultracentrifuge tube, and the DNA was prepared by treatment with Proteinase K and SDS, followed by repeated extractions with phenol-chloroform and ethanol precipitation. The DNA content and purity were observed by spectrophotometry and by agarose gel electrophoresis after restriction enzyme digestions.

### Cell culture

Vero cells (ATCC; Manassas, VA) were propagated in M199 medium supplemented with 5% fetal bovine serum and gentamycin. HaCaT cells (Department of Dentistry, University of Turku; (36)) were propagated in DMEM medium with HEPES buffer, supplemented with 7% fetal bovine serum and gentamycin. SH-SY5Y neuroblastoma cells (K. Åkerman, Åbo Akademi University, Turku, Finland) were propagated in DMEM (high glucose) medium supplemented with 10% fetal bovine serum, 2 mM L-glutamine and gentamycin. Initial differentiation of SH-SY5Y cells involved culture for 10 days in DMEM/F-12 medium containing 5% fetal bovine serum, 10 µM All-trans retinoic acid (Sigma), 2 mM L-glutamine and gentamycin. Thereafter SH-SY5Y cells were transferred on Matrigel-coated (B-D) 96-well plates and the medium was changed to serum-free DMEM/F-12 medium containing 10 µM all-trans retinoic acid (Sigma), 0.5 µg/ml of brain-derived neurotrophic factor (Millipore), 2 mM L-glutamine and gentamycin.

### Acyclovir resistance testing

The sensitivity of each HSV strain to acyclovir was tested in a microplate format. Vero cells grown on 96-well cell culture plates were treated with cell culture medium (DMEM with 5% fetal bovine serum) supplemented with acyclovir (ACV; Sigma), in concentrations of 128 µg/ml to 0.03125 µg/ml (1:4 serial dilutions). Duplicate wells containing each ACV dilution, and wells without ACV, were infected with 100 PFU of each virus. As a control, the HSV-1 delta305 virus was included; it is resistant to ACV due to deletion of its thymidine kinase gene (37). Infected cells were incubated at +37°C, at 5% CO_2_, for 72 hours before fixation with methanol and staining with crystal violet. The reduction of plaque numbers at each ACV dilution was observed in comparison with wells infected without ACV. A logistic fit curve was used for determination of IC_50_ values. A virus with an IC_50_ value of over 2 µg/ml (of ACV) was considered resistant (38).

### Image acquisition

Photomicrographic images of viral plaques were obtained using a Zeiss Primovert inverted microscope with Plan-Achromat 4x and 10x objectives, recorded using an AxioCam ERc 5s camera, and analyzed using Zeiss ZEN 2012 software.

### Next generation sequencing

Viral nucleocapsid DNA was sheared on a Covaris M220 (parameters: 60 seconds duration, peak power 50, 10% duty cycle, 4C). We used the Illumina TruSeq DNA Sample Prep kit to prepare barcoded sequencing libraries, according to the manufacturer’s protocol for low-throughput sample handling. Libraries were quantified and assessed by Qubit (Invitrogen, CA), Bioanalyzer (Agilent), and library adapter qPCR (KAPA Biosystems). Illumina MiSeq paired-end sequencing (2 x 300 bp) was completed according to manufacturer’s recommendations, using 17 pM library concentration.

A consensus viral genome for each strain was assembled using a *de novo* viral genome assembly (VirGA) workflow (10). This approach begins with quality control, including removal of contaminating host sequences, adapters from library preparation, and imaging artifacts. Next VirGA iterates through multiple *de novo* assemblies using SSAKE, which we then combine into longer blocks of sequence (contigs) using Celera and GapFiller. VirGA uses Mugsy alignment to match these contigs to the HSV-1 reference genome (strain 17, GenBank accession JN555585). The best-matched contigs are stitched into a single consensus genome, which is then annotated and subjected to additional quality control measures. These include an examination of coverage (sequence read) depth, detection of minority variants within each consensus genome, and manual inspection of gaps and low coverage areas. See **Table 2** for GenBank accessions.

**Table 2.**
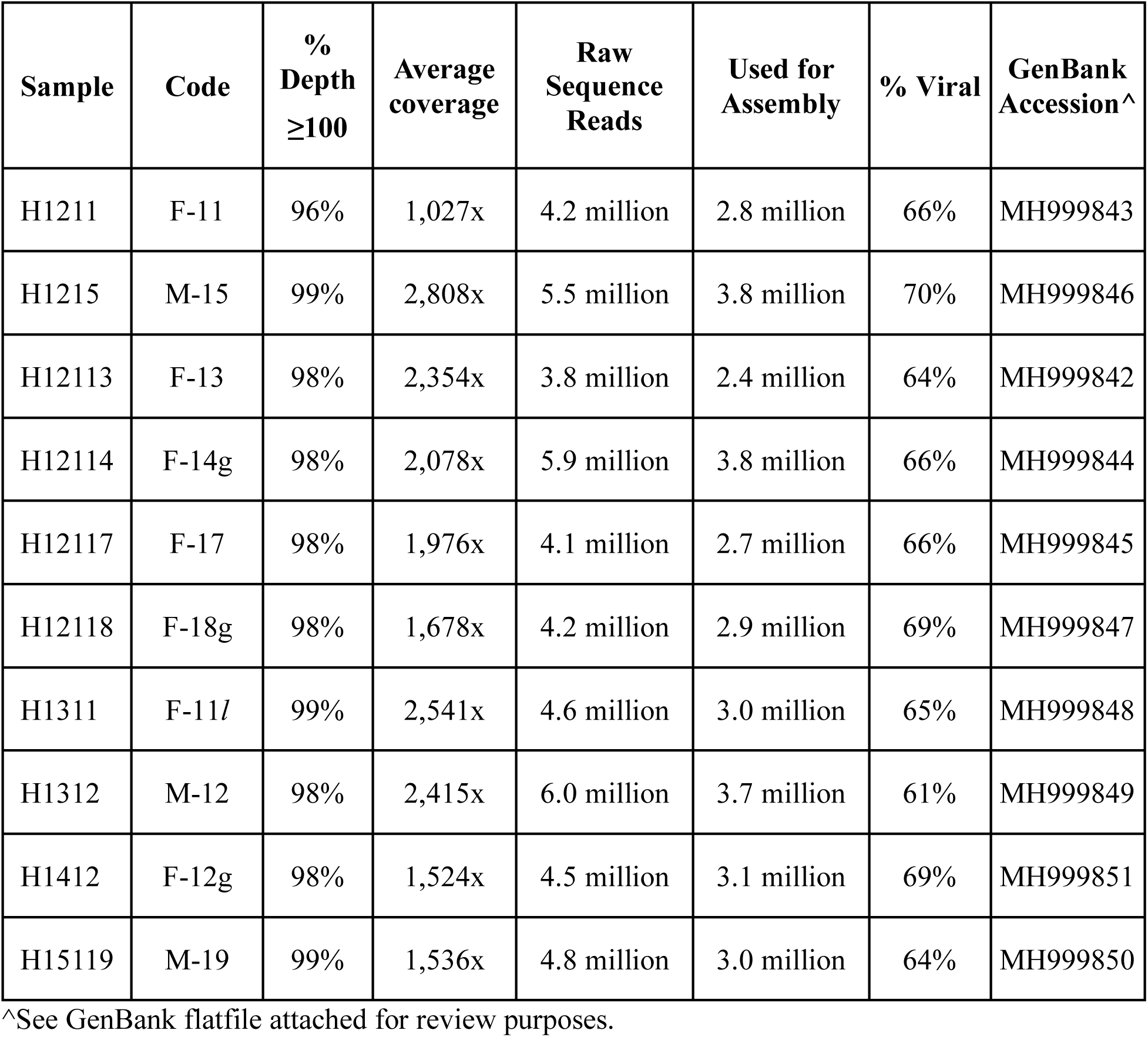
Sequencing statistics for Finnish HSV-1 strains.

| **Sample** | **Code** | **%**<br>**Depth**<br>**≥100** | **Average**<br>**coverage** | **Raw**<br>**Sequence**<br>**Reads** | **Used for**<br>**Assembly** | **% Viral** | **GenBank**<br>**Accession<sup>^</sup>** |
| --- | --- | --- | --- | --- | --- | --- | --- |
| H1211 | F-11 | 96% | 1,027x | 4.2 million | 2.8 million | 66% | MH999843 |
| H1215 | M-15 | 99% | 2,808x | 5.5 million | 3.8 million | 70% | MH999846 |
| H12113 | F-13 | 98% | 2,354x | 3.8 million | 2.4 million | 64% | MH999842 |
| H12114 | F-14g | 98% | 2,078x | 5.9 million | 3.8 million | 66% | MH999844 |
| H12117 | F-17 | 98% | 1,976x | 4.1 million | 2.7 million | 66% | MH999845 |
| H12118 | F-18g | 98% | 1,678x | 4.2 million | 2.9 million | 69% | MH999847 |
| H1311 | F-11l | 99% | 2,541x | 4.6 million | 3.0 million | 65% | MH999848 |
| H1312 | M-12 | 98% | 2,415x | 6.0 million | 3.7 million | 61% | MH999849 |
| H1412 | F-12g | 98% | 1,524x | 4.5 million | 3.1 million | 69% | MH999851 |
| H15119 | M-19 | 99% | 1,536x | 4.8 million | 3.0 million | 64% | MH999850 |

<sup>^</sup>See GenBank flatfile attached for review purposes.

### Intra-strain minority-variant detection

Minority-variant (MV) positions within each *de novo* assembled genome were determined using VarScan v2.2.11 as previously described (19, 39). Conservative variant calling parameters to eliminate sequencing-induced errors were set at: minimum allele frequency ≥0.02; base call quality ≥20; read depth at the position ≥100; independent reads supporting minor allele ≥5. MVs containing ≥90% unidirectional strand support were excluded from further analyses (40, 41), as were those occurring within 10 base pairs of one another. MVs passing quality control were mapped to respective genomes and assessed for mutational effect using SnpEff and SnpSift (42, 43).

### Phylogenetic and recombination analyses

DNA sequences were aligned using the Kalign algorithm included in eBioTools. To avoid inferences caused by false phylogenetic signals, all gap and repeat regions were excluded prior to this analysis. Repeat regions not leading to gaps were also excluded since these may contain single nucleotide differences that have been shuffled into new positions by random aspects of tandem-repeat alignment. Furthermore, nucleotides that were marked as “N” in GenBank-derived strains were excluded, along with the corresponding aligned nucleotides in remaining sequences. Complete genomes in GenBank harbouring long stretches of “N” were not included in the analysis (e.g. strain B3x_1_5). The network is based on an alignment of 116,610 nucleotides, which is more than 75% of the entire HSV-1 genome.

The phylogenetic network was constructed by using SplitsTree4. It is a NeighborNet with OrdinaryLeastSquares Variance depicted as a RootedEqualAngle splitstree. A recombination test on the complete dataset was performed by using the Phi test for recombination implemented in SplitsTree4. Bootstrapping values greater than 80 are shown for clades with three or more strains.

### Restriction fragment length polymorphism (RFLP) analysis

A cytoplasmic viral DNA preparation was modified from the protocol described by Igarashi et al (44). Subconfluent Vero cell cultures were infected at an m.o.i. of 0.1, and incubated at +34°C for 2 days until the cytopathic effect was complete. The cells were collected in 150 mM NaCl, 10 mM Tris pH 7.6, 1.5 mM MgCl buffer, with 0.1% Nonidet P-40. The nuclei were pelleted, and DNA was extracted from the supernatant by two successive extractions with phenol-chloroform. DNA was recovered by ethanol precipitation. 12.5 µg of each viral DNA was diluted in 15 µl solution containing sterile H_2_O, FastDigest 10X green buffer and FastDigest BamHI or SalI restriction enzyme (Thermo Scientific). Electrophoresis was run in 0.8 % TBE-agarose gels, with GeneRuler mix DNA ladder, for 24 h at 45 V before imaging. In order to separate large DNA fragments after SalI digestion, electrophoresis was continued for additional 36 h at 30 V.

## Results

### Comparison of growth properties of Finnish HSV-1 strains in vitro

A random set of ten circulating Finnish HSV-1 isolates were selected from residual laboratory diagnostic specimens (**Table 1)**. We first examined *in vitro* phenotypic characteristics of these HSV-1 isolates, by comparing their overall titer and rate of intracellular virus vs. extracellular (released) virus production at 24 hours post infection (hpi) of a range of cell types (**Figure 1**). These included non-human primate kidney-derived epithelial (Vero) cells (**Figure 1A**), human keratinocyte (HaCaT) cells (**Figure 1B**), and human neuroblastoma (SH-SY5Y) cells in a mixed, undifferentiated state (**Figure 1C**), as well as a differentiated, neuronal state (**Figure 1D**). Compared to epithelial and keratinocyte cells, overall viral production was markedly reduced in neuronal precursor cells and differentiated neuronal cells (compare **Figure 1AB** vs. **CD**). The wild-type HSV-1 reference strain 17+ replicated to significantly higher titers than any of the circulating clinical isolates in Vero cells, which are standardly used for HSV propagation, and to a lesser extent strain 17+ replicated at higher titer in keratinocytes as well (**Figure 1**; p<0.05, 10-100 fold higher viral amounts). Different clinical isolates excelled at producing virions in each cell type, with no clear patterns of most- or least-efficient viral production or release across all cell types. The amount of extracellular released virus was less than 10% of the total virion production, in all cell types except the undifferentiated neuronal precursor cells (**Figure 1, insets**). Each virus strain was also tested for acyclovir (ACV) resistance. Despite some variation in ACV susceptibility (IC50 values), none of the circulating Finnish viruses was considered resistant to ACV (**Table 1**). There were also no significant differences among the 10 clinical isolates in the plaque morphology or the type of cytopathic effect induced in Vero cell cultures (data not shown).

**Figure 1:**
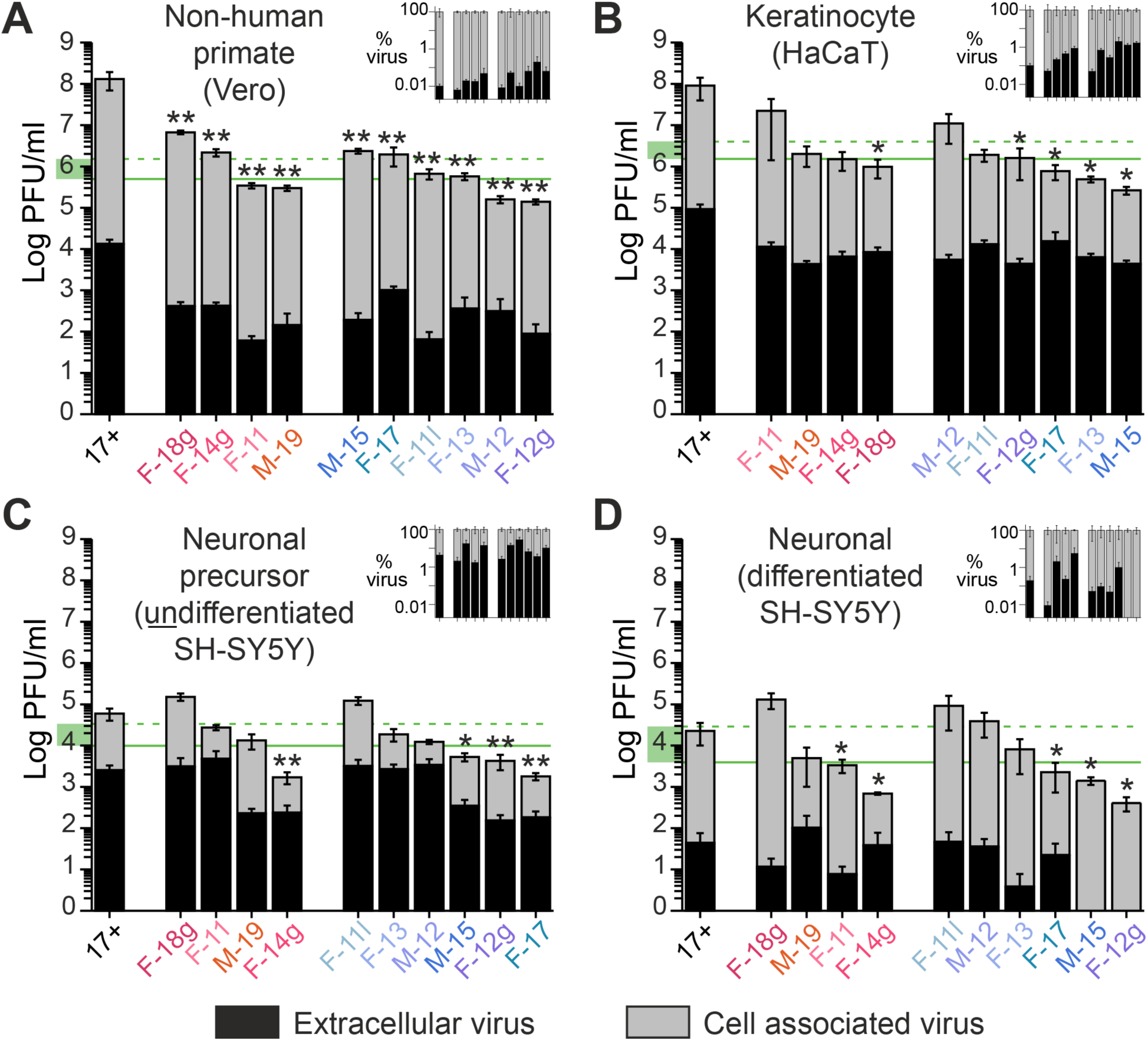
Comparison of growth properties of Finnish HSV-1 strains reveals the uncoupling of phenotypic and geographic variation. Histograms comparing virus production for ten Finnish strains of HSV-1, as compared to the HSV-1 reference strain 17+, in different cell types. Panel **(A,** non-human primate) shows Vero monkey kidney cells, which are defective in interferon signaling; **(B)** shows HaCaT human keratinocytes; **(C)** shows undifferentiated human SH-SY5Y neuroblastoma cells (neuronal precursor cells); **(D)** shows differentiated human SH-SY5Y neuronal-like cells. Viruses are plotted in descending order based on their viral production. For ease of comparison, the strains are color-coded and divided into the genomic sub-groups used in later figures (see **Figure 3** for illustration). Each histogram bar plots infectious virions in the cell-associated fraction (gray) vs. those released from cells (black). A horizontal line shows the median (solid green) and the average (dashed green) amount of virus production for the ten Finnish HSV-1 strains in each cell type, with a highlighted box (green) on this y-axis added to highlight the multi-log decrease in viral production in **C-D** vs. **A-B**. Each panel has a smaller graph (**inset**) where the extracellular released virus (black) is plotted as a percentage of total virus production (gray). Viruses in each inset are plotted in the same order as the matched panel graph. Bar shows SEM; * = p=<0.05, ** = p<0.001 as compared to the total virus amount (cell associated + extracellular), for each clinical isolate vs HSV-1(17+) reference strain.

### Genetic and genomic analysis of Finnish HSV-1 strains

Based on prior data suggesting the effects of successive waves of migration on the human population in Finland (45, 46), we next assessed the overall genetic diversity of these ten HSV-1 isolates. We first performed a restriction fragment length polymorphism (RFLP) analysis on viral genomic DNA, which revealed at least two broad patterns of variation (**Figure 2**). The diversity of bands led us to examine these genetic differences with greater precision using high-throughput, deep genome sequencing (HTS) and comparative genomics analysis (see **Methods** for details). A previously described bioinformatics pipeline was used to *de novo* assemble a full-length consensus genome for each strain **(Table 2)**. A viral consensus genome represents the most common nucleotide detected at each nucleotide position in that viral population. All viral genomes had an average coverage depth between 1,000-2,800-fold, with >96% of the genome covered at a depth exceeding 100-fold **(Table 2)**. As observed in prior publications, the few areas with <100X coverage correlated with regions of high G+C-content, and/or highly repetitive sequences (47, 9, 10). The >1,000X average coverage depth enabled us to compare the full genetic complement of the viral genome for each strain, to analyze differences both between and within each viral strain population, and to compare them to previously described viral genomes.

**Figure 2:**
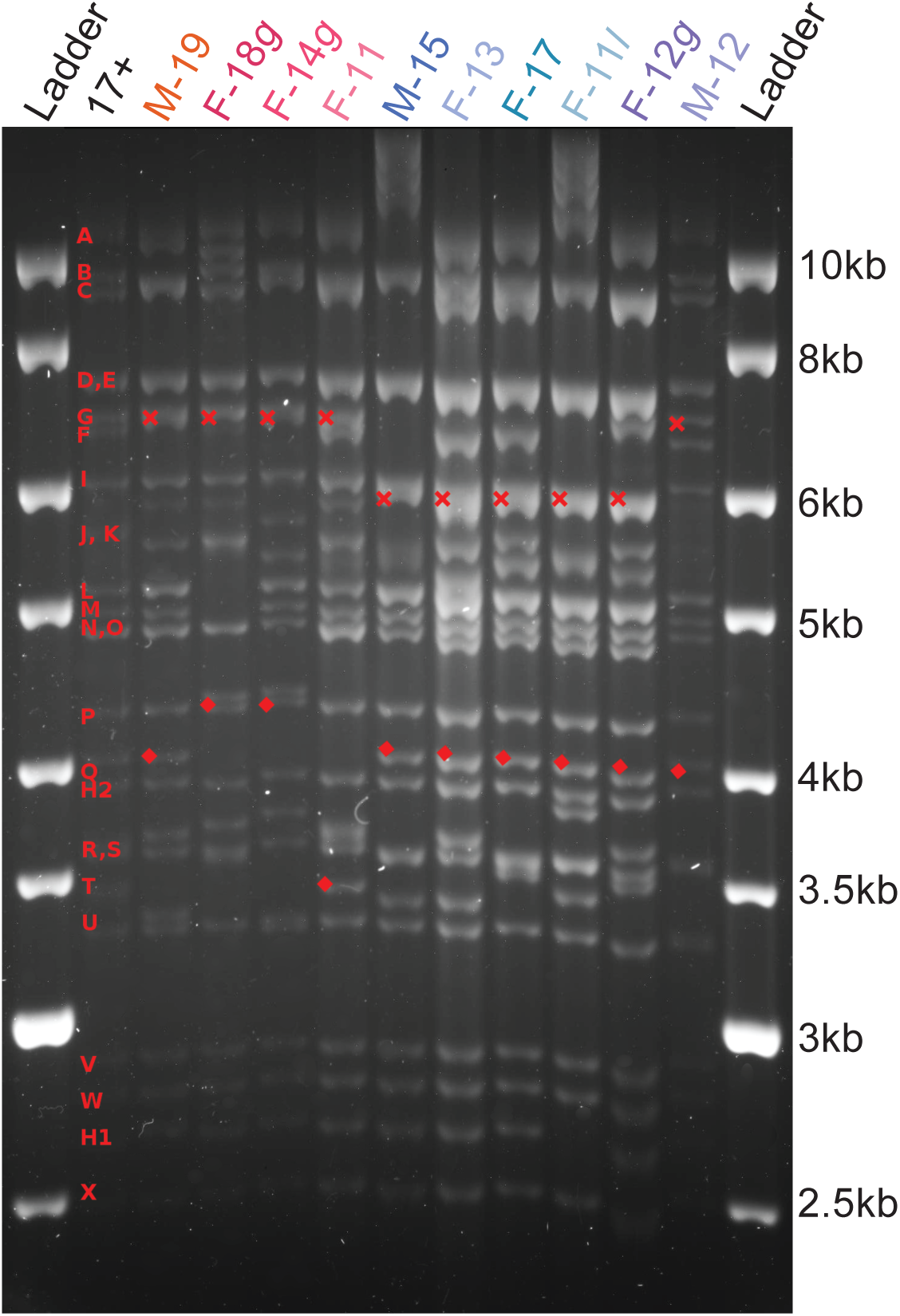
RFLP comparison reveals at least two sub-groups of Finnish HSV-1 strains. Viral genomic DNA was digested with SalI and separated via electrophoresis to distinguish overall genetic patterns. Classically defined SalI fragment notations (red alphabet letters) are indicated on the left (54). Red crosses (x) indicate variation in the size of SalI fragment G, while diamonds indicate variation in the SalI Q fragment. For ease of comparison, the strains are color-coded the same as in later figures (see **Figure 3** for illustration).

### Genetic relatedness of Finland HSV-1 samples

First, we used the viral consensus genomes to compare how overall variation in these ten Finnish HSV-1 genomes related to the patterns observed by RFLP analysis (**Figure 2**). Whole genome alignments were created using the ten Finnish HSV-1 genomes, as well as these ten in conjunction with a diverse set of previously published HSV-1 genomes (see Methods and **Supplemental Table S1** for a full list). We then used SplitsTree to create a phylogenetic network that revealed the relatedness of the ten Finnish viral genomes to each other (**Figure 3A**) and to previously described HSV-1 genomes (**Figure 3B**). This approach revealed that the Finland HSV-1 genomes separated into two sub-groups, which appear to relate to previously recognized Asian and European / North American clades. These data echo those of the initial RFLP analysis.

**Figure 3.**
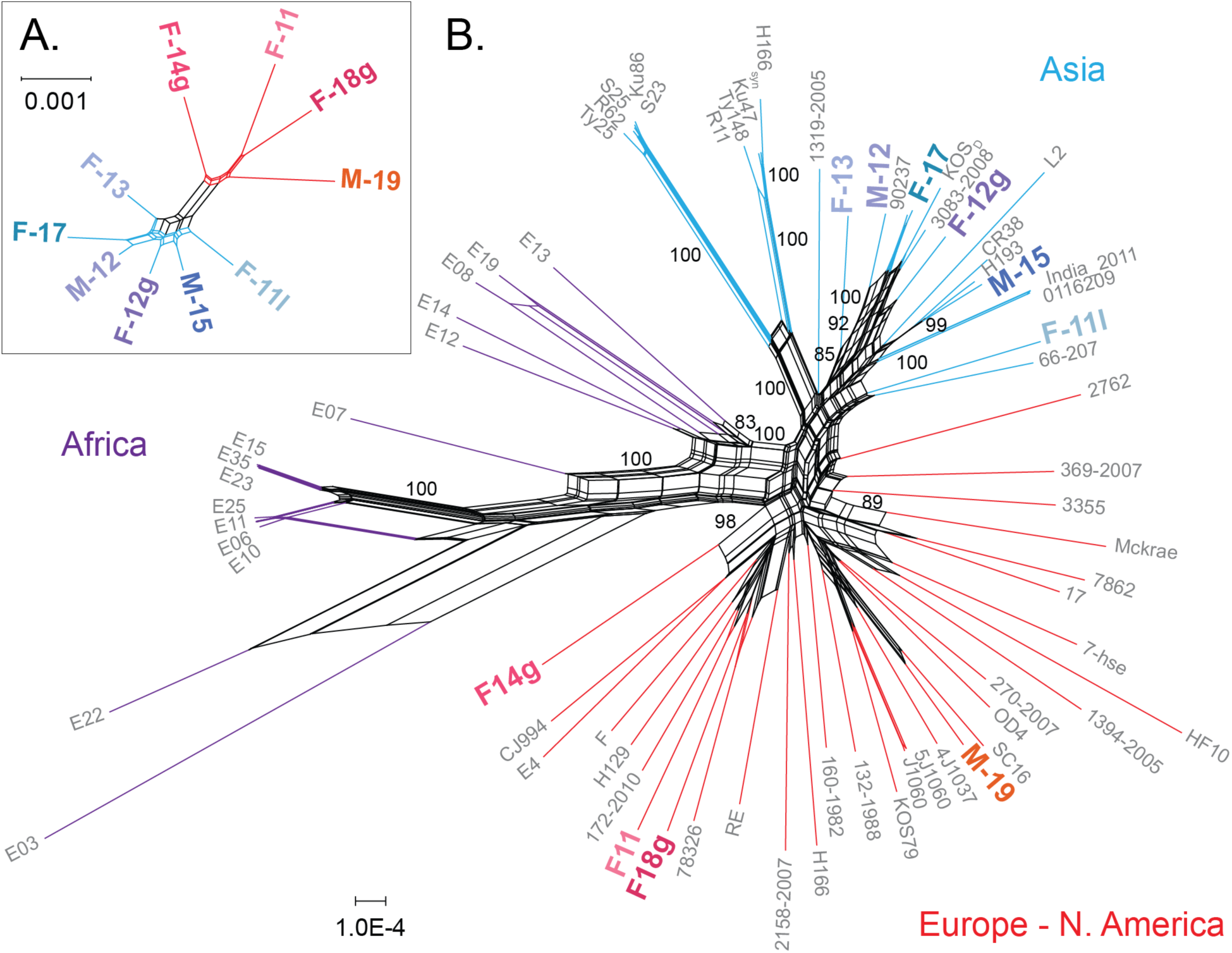
A phylogenetic network of genetic relatedness reveals that Finland HSV-1 isolates separate into two sub-groups. A phylogenetic network of genetic relatedness was constructed using SplitsTree4 to demonstrate how the ten new Finland HSV-1 isolates relate to each other (**A**) and to previously described HSV-1 genomes from a diversity of locations (**B**). Geographic clustering of branches and clades resemble those previously described (9, 16–18) and are colored by origin of strains: red for Europe and North America, blue for Asia, and purple for Africa. Finland HSV-1 genomes are indicated in bold, and separate into two sub-groups (**A**), which appear to relate previously recognized Asian and European and North American clades (**B**). Bootstrap values larger than 80 supporting clades with three or more strains are shown on the network. The extent of recombination among HSV-1 strains is demonstrated by the parallel edges in the network and a phi-test for recombination. The isolate name, country of origin, GenBank accession, and reference(s) for each previously described HSV-1 genome are listed in **Supplemental Table S1**.

### Protein-coding variation in Finland HSV-1 samples

Next, we examined more fine-scale genetic differences in these HSV-1 isolates, which are invisible at the level of RFLP analysis. Here we examined DNA- and amino acid (AA)-alignments for each protein-coding region of the HSV-1 genome (**Supplemental Table S2-S3**). We found that from zero to 8% of each coding region differed at the AA level between the ten Finnish viral genomes (**Figure 4**). Only a handful of small coding regions (UL20, VP26 (UL35), UL49A, UL55) showed no AA variation at all between viral isolates. Consistent with previous findings (9), gL (UL1), UL11, UL43, gG (US4), and gJ (US5) were among the most divergent proteins. Together these inter-strain genetic differences in 70 HSV-1 proteins in **Figure 4** provide ample opportunities to generate the phenotypic diversity observed in **Figure 1**. Overall, these levels of AA coding diversity reflect those seen in previous analyses (9), and between other known HSV-1 genomes.

**Figure 4.**
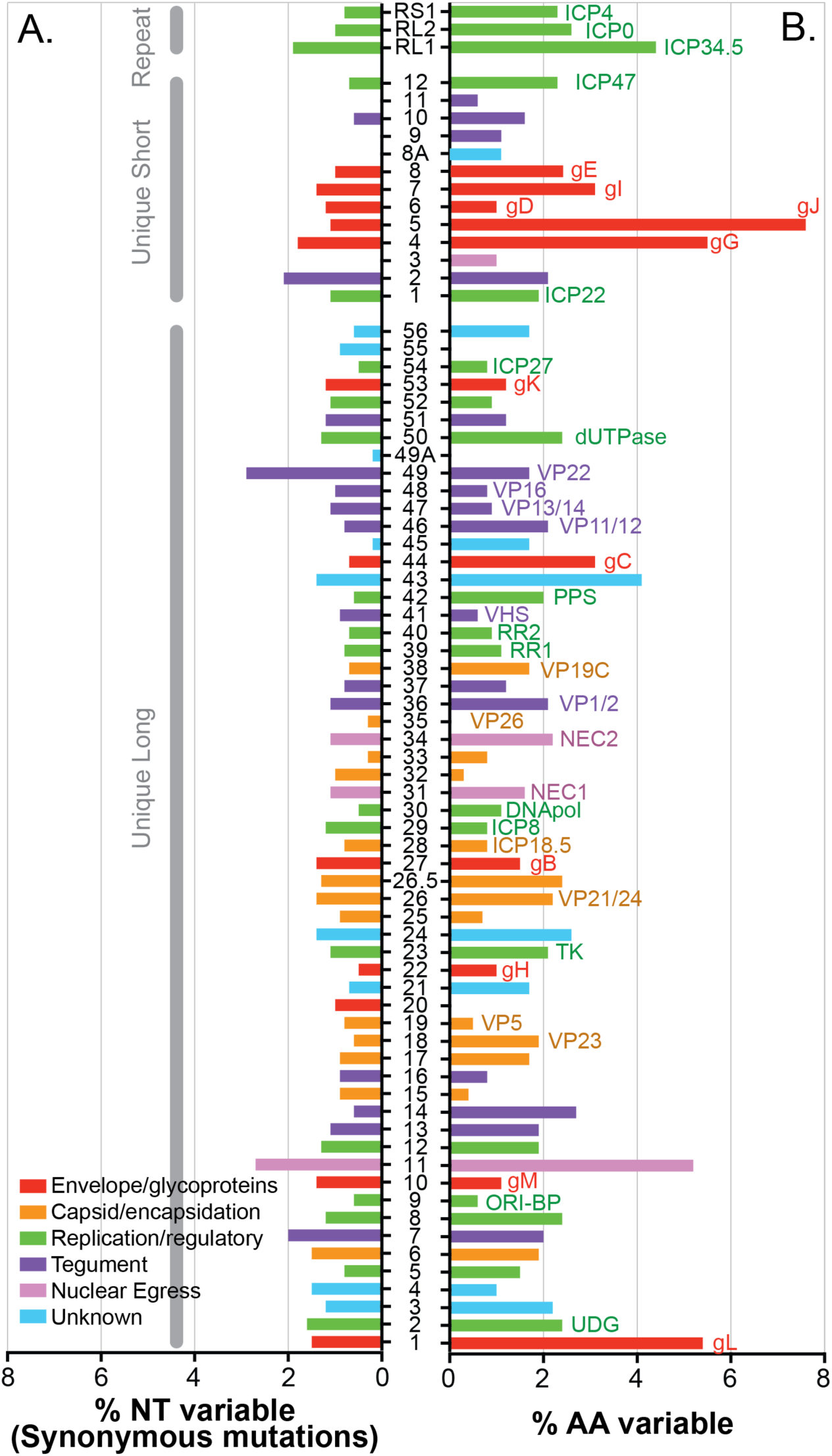
Substantial protein-coding variation exists between Finland HSV-1 isolates. Histogram depicts the percent of synonymous nucleotide differences per gene (**A**) and the percent amino acid (AA) variation per protein (**B**) for the ten Finnish HSV-1 strains compared here. Labels on the left in (**A**) indicate the repeat, unique short (US) and unique long (UL) regions that contain each of the sequentially-numbered viral coding regions. Where possible, the common name of each protein is shown to the right of the appropriate histogram bar in (**B**), e.g. UL1 is also known as gL. Nucleotide and AA differences were quantified at the consensus level of each viral genome. See **Supplemental Table S2** for additional data on the number of nucleotide differences and dN/dS ratio for each gene.

### Minority allelic variants in Finland HSV-1 viral populations

Each viral consensus genome represents the most common nucleotide detected at each position in the viral genome. In contrast, minority variants (MV) within each viral population are rare alleles, presented in <50% of the sequenced reads. These MV may rise to greater prevalence during viral spread to new niches or hosts, or under selective evolutionary pressure. For each viral genome, we examined the number of MVs detected in each genome. Stringent quality-control criteria were used to reduce the number of false-positive MVs (see **Methods**), and only those MV detected at ≥2% prevalence were used in this analysis. MVs were found dispersed across each genome (**Figure 5A**), mostly in intergenic regions, and were mostly of low frequency (**Figure 5B**). A concentration of MVs was found in the internal-repeat region, which may result from stochasticity in tandem repeat alignment in this area, and/or from the reduced selective pressures on intergenic sequences found in this region. The MVs detected in US7 (gI) occur at the same tandem-repeat site as the consensus-level AA variations in this gene (**Figure 6**). Taken together, these results define the level of inter-host (**Figure 4**) and intra-host (**Figure 5**) variation seen in the Finnish HSV-1 strains described herein.

**Figure 5.**
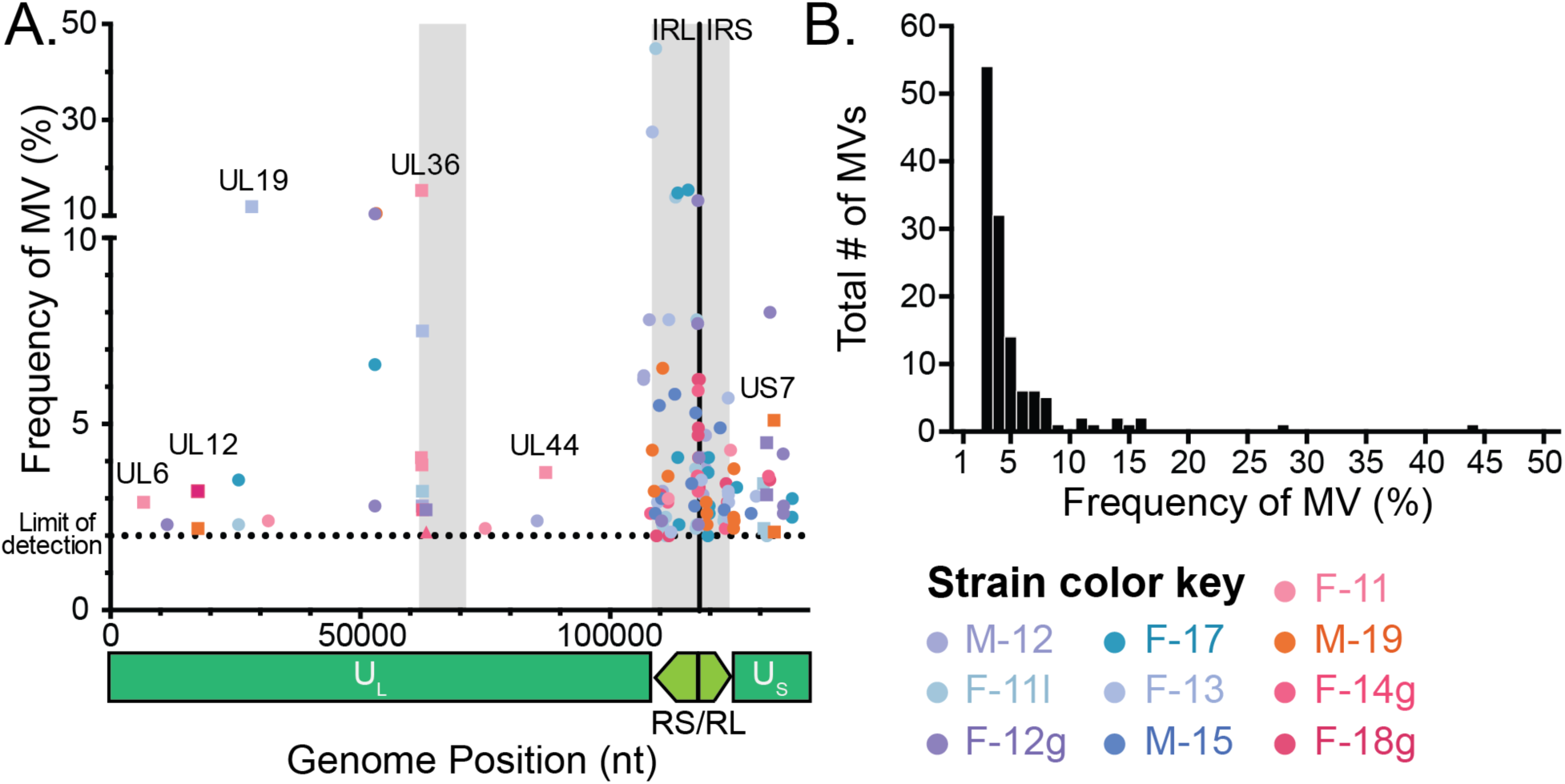
Scatter plot illustrates the location and low frequency of minor variants in each Finnish HSV-1 genome. **(A)** Scatter plot demonstrates the position (x-axis) and frequency (y-axis) of minority variants detected in each Finnish HSV-1 genome. Spheres on the graph indicate minor variants in intergenic regions. Squares with gene names indicate MVs located in genes. A color key consistent with that used other figures is included at the lower right. A genome diagram is shown below the x-axis for reference. Areas containing large numbers of tandem repeats, e.g. around UL36 in the UL region and throughout the RS/RL region, are denoted as gray blocks on the graph. The limit of detection was set at 2% (see Methods for details). **(B)** Histogram depicts the total number of minority variants (y-axis) in each frequency bin (x-axis), summed across all ten Finnish HSV-1 strains. Bins are in 1% increments.

**Figure 6.**
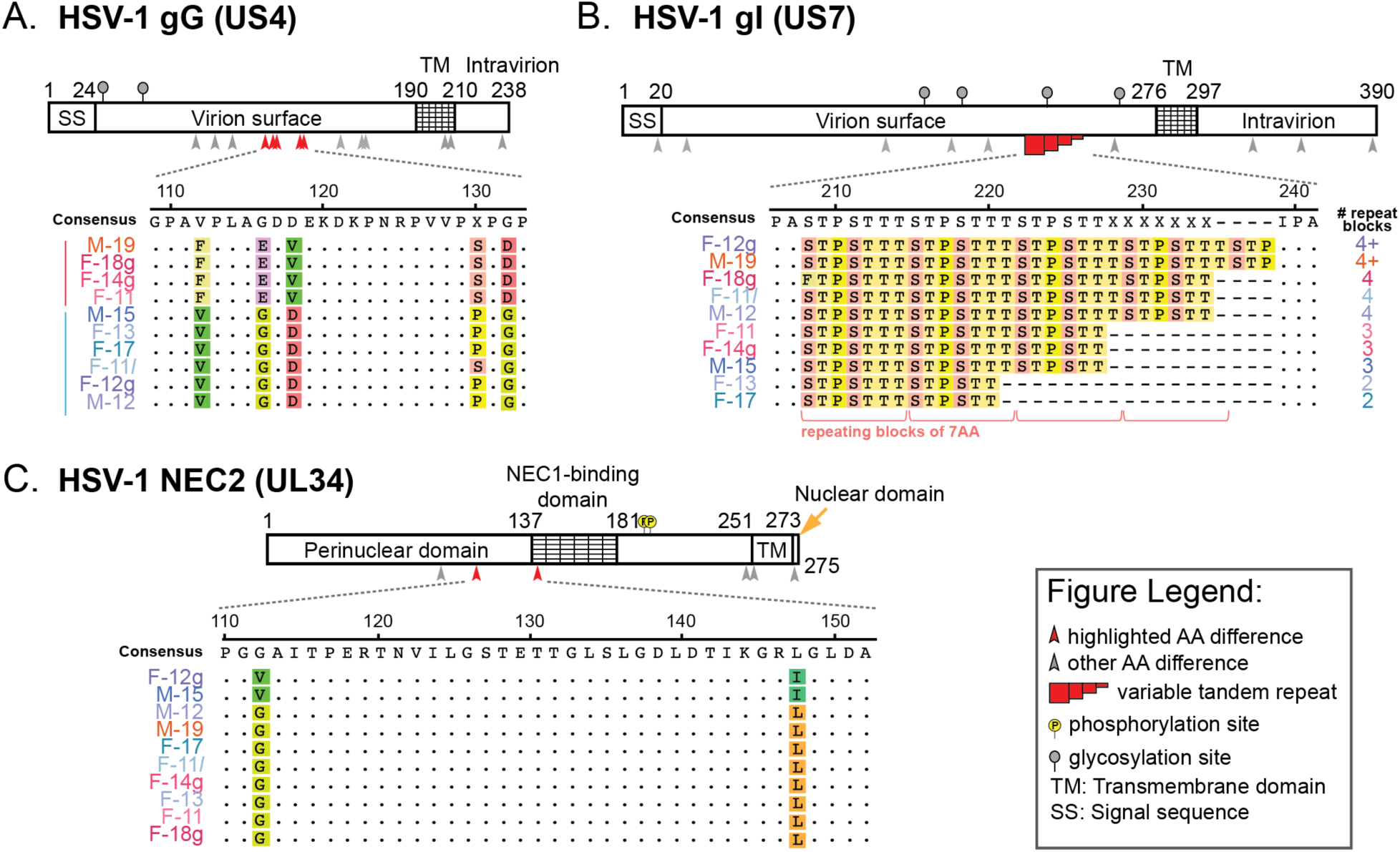
Variations in individual genes provide a better insight into Finnish HSV-1 strain sub-groups and of specific cellular phenotypes. Examples are shown here of variation in individual HSV-1 genes that reflect either the Finnish HSV-1 strain sub-groups, or that correlate with and potentially influence specific cellular phenotypes. Diagrams depict three examples of HSV-1 proteins with known functional domains and post-translational modifications (e.g. phosphorylation, glycosylation; see Legend). A subset of each protein is highlighted via an AA alignment of the ten clinical HSV-1 isolates. (**A**) Coding differences in the secreted and virion surface glycoprotein G (gG; US4) correlate with the overall genomic sub-groups (clades) shown in **Figure 3**. (**B**) Variation in copy number of a seven AA tandem repeat block in glycoprotein I (gI; US7) changes the length of a repeating tract of mucin-type O-linked glycosylation sites, as described previously in clinical isolates of HSV-1 (16, 48).(**C**) AA differences in the nuclear egress protein NEC2 (UL34) were observed in the two strains (M-15, F-12g) which showed no detectable extracellular virion release in differentiated neuronal cells (**Figure 1D**). See Discussion for additional insights on the potential phenotypic impacts of the variations shown in B-C.

### Patterns of variation in Finnish HSV-1 strains

Finally, we considered whether any of the observed patterns of genomic or phenotypic variation could be linked to fine-scale coding variations in these ten Finnish HSV-1 isolates. The genomic sub-groups detected by RFLP (**Figure 2**) and phylogenetic network analysis (**Figure 3**) were reflected in several coding variations that correlated with phylogenetic group. For example, coding differences in the secreted and virion surface glycoprotein G (gG; US4) correlated with the two Finnish phylogenetic sub-groups (**Figure 6A**; see **Figure 3** for phylogenetic sub-groups).

Other impacts on coding variation may result from changes in copy number at tandem repeats. Variation in tandem repeat length in glycoprotein I (gI; US7) leads to changes in the length of a repeating tract of mucin-type O-linked glycosylation sites (**Figure 6B)**, akin to that previously described in clinical isolates of HSV-1 (16, 48). Finally, there are detectable patterns of AA variation that correlate with phenotypic differences in viral fitness in specific cell types. Coding differences in the nuclear egress complex protein NEC2 (UL34) (**Figure 6C**) were observed in the two strains (F-12g, M-15) which showed no detectable extracellular virion release in differentiated neuronal cells (**Figure 1D**). The terminally differentiated and non-dividing state of these neuronal cells may constitute a sensitized background to detect impacts on virion egress. Taken together, these examples illustrate the type and degree of variation present in these strains, and highlight potential genetic insights into viral phenotype.

## Discussion

In this study, we described the first-ever HSV-1 genomes from Finland. Two distinct sub-groups or clades were observed in the ten Finnish clinical isolates, with four strains clustering in one clade and six in another. These groups were detectable using both classical RFLP approaches (**Figure 2**) and deep-sequencing methods for whole-genome analyses (**Figure 3**). These data appear to corroborate with previous findings on colonization of Finland, with migration from Europe in the south/southwest and from Asia in the northeast (26–28). However, we found that phenotypic differences between these isolates were uncoupled from the overall genomic patterns that grouped them into two geographic clades. While plaque morphology was consistent across isolates (data not shown), the amount of virus production, extracellular virion release, and acyclovir sensitivity differed between isolates and across cell types (**Figure 1** and **Table 1**). We detected gene-specific patterns of genetic variation that may impact protein function. Future studies will need to dissect individual genetic variations in each viral isolate in order to test their precise impact(s) on viral fitness.

When analyzing the replication capabilities of these recently isolated clinical samples in multiple cell types, we found that they replicated to lower overall titers and produced fewer extracellular virions than the lab-adapted HSV-1 strain 17+, regardless of cell type (**Figure 1**). This difference was more pronounced in non-human primate epithelial and human keratinocyte cells, and was more subtle in neuronal precursors and differentiated neuronal cells. There did not appear to be any clear differences in titer or extracellular virus production between geographic clades. There also seemed to be no noticeable differences in plaque morphology among these samples, with plaque size and cellular cytopathic effect being similar across all samples (data not shown). Although we report findings in a number of cell lines in this study, it will be important for future studies to examine each virus and cell-type pairing separately, since these data demonstrate that maximal viral fitness in one cell type is not generally predictive of fitness in another.

We observed several potential connections between genetic differences and phenotypic differences. For instance, we detected an AA difference in the UL34 (NEC2) binding domain in F-12g and M-15 **(Figure 6C**). Previous work has shown that the viral proteins UL31 (NEC1) and UL34 (NEC2) form a nuclear egress complex that localizes to the perinuclear space and plays a critical role in egress of viral capsids from the nucleus (49, 50). In these two clinical isolates (F-12g and M-15), there was no measurable amount of extracellular virus produced in differentiated neuronal cells (**Figure 1D**). Further studies will be needed to examine if this phenotype is linked to a disruption of NEC1 and NEC2 binding and impaired nuclear egress for isolates F-12g and M-15. Variation in the copy-number of tandem repeats is yet another way that HSV-1 isolates can generate genetic and potential phenotypic variation (9, 16, 51–53). Glycoprotein I (gI; US7) contains a mucin-like domain with repeating units of serine, threonine, and proline (**Figure 6B**). This domain varies in length across these isolates. This domain has been previously shown to serve as a site of O-linked glycosylation, with longer tracts (e.g. 8 repeating blocks of 7 AA each) being more heavily glycosylated than short tracts (e.g. 2 repeating blocks) (16, 48). A prior study of over 80 Swedish HSV-1 isolates found roughly equal distributions of isolates with two, three, or four repeating blocks in this mucin-like gI domain (16, 48). Half of the Finland HSV-1 isolates analyzed here have at least four repeating units, suggesting the potential for gI to be more heavily glycosylated in these isolates. These data generate hypotheses for future investigations of genotype-phenotype links in these and other new clinical isolates.

This study provides one of the first combinations of phenotypic analyses that span multiple cell types, alongside full comparative genomic analyses that span both phylogenetic and gene-specific analyses. One comparison not yet explored in this or prior studies is the specific pairing of each virus with cells derived from the human source of each isolate. This approach is now technically feasible, using induced pluripotent stem cell technology to generate specific cell types from a single source such as a buccal swab. The rich genealogical databases in Finland suggest an opportunity to not only pair these analyses at the human-cell and virus level, but also to link observed phenotypes of cellular infection to each patient’s clinical, genetic, and/or familial history of herpesvirus disease. Future studies may also seek to link human interaction data, for instance contact networks and/or travel history, to infer how human movement and interactions patterns influence pathogen spread and population dynamics. We anticipate that in the future, this approach would yield fruitful insights not only into direct clinical outcomes of herpesvirus disease, but also generate hypotheses about herpesvirus co-morbidities.

## Acknowledgements

The authors thank professor Päivi Onkamo, and the members of the Hukkanen and Szpara labs for helpful feedback and discussion. We acknowledge Ritva Kajander for preparation of the viral stocks and the viral nucleocapsid DNA, and BSc Kiira Kalke and BSc Mira Hörkkö for help in imaging the viral plaques. This work was supported by research startup funds (MLS) from the Pennsylvania State University, and by a grant from the Pennsylvania Department of Health using CURE funds (MLS). The Department specifically disclaims responsibility for any analyses, interpretations, or conclusions. The funding from the Jane and Aatos Erkko Foundation grant #170046 is acknowledged. We thank Prof. Bernard Roizman, University of Chicago, for the HSV-1 delta305 virus.

## Supplemental Tables

**Supplemental Table S1:**
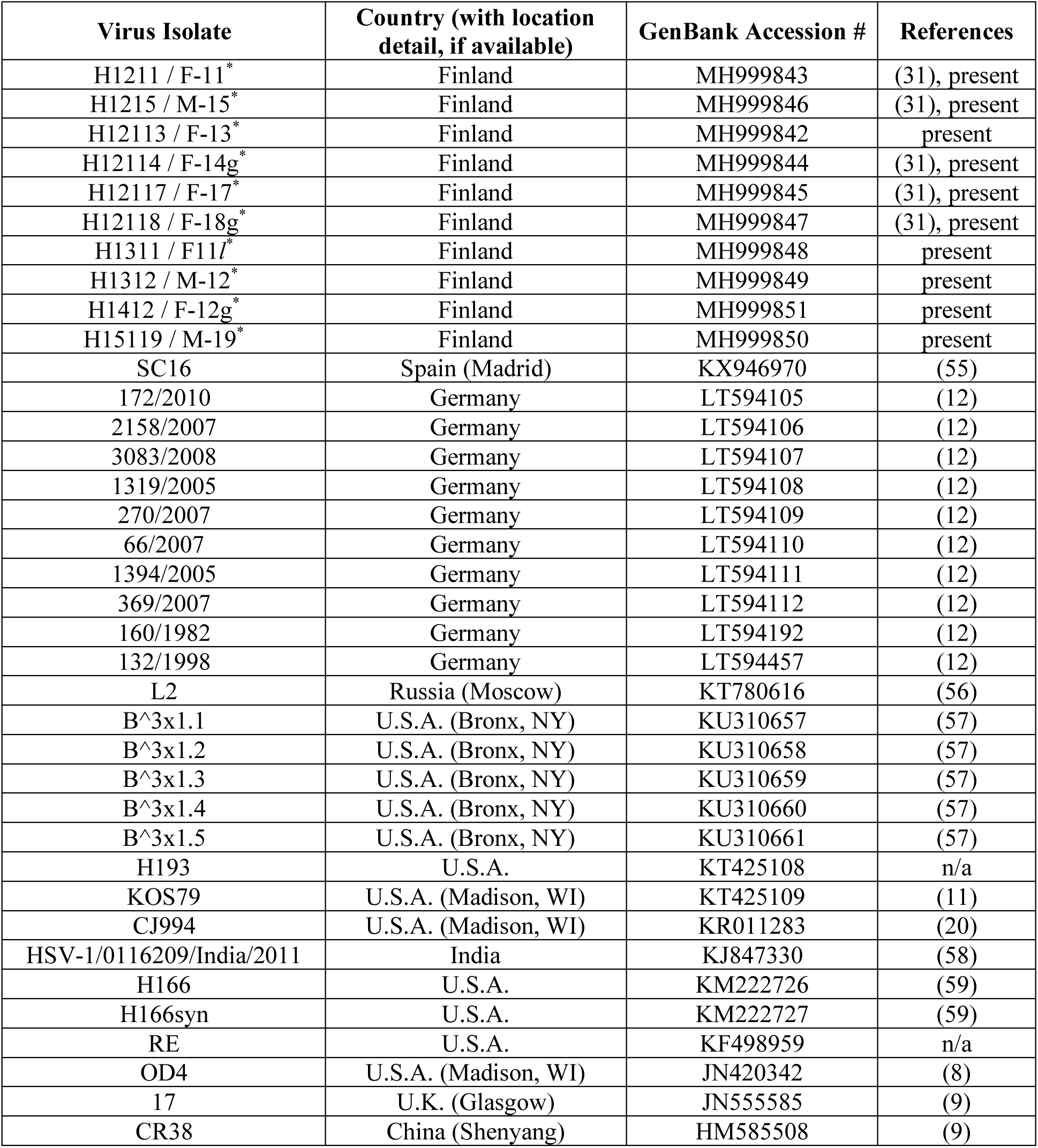
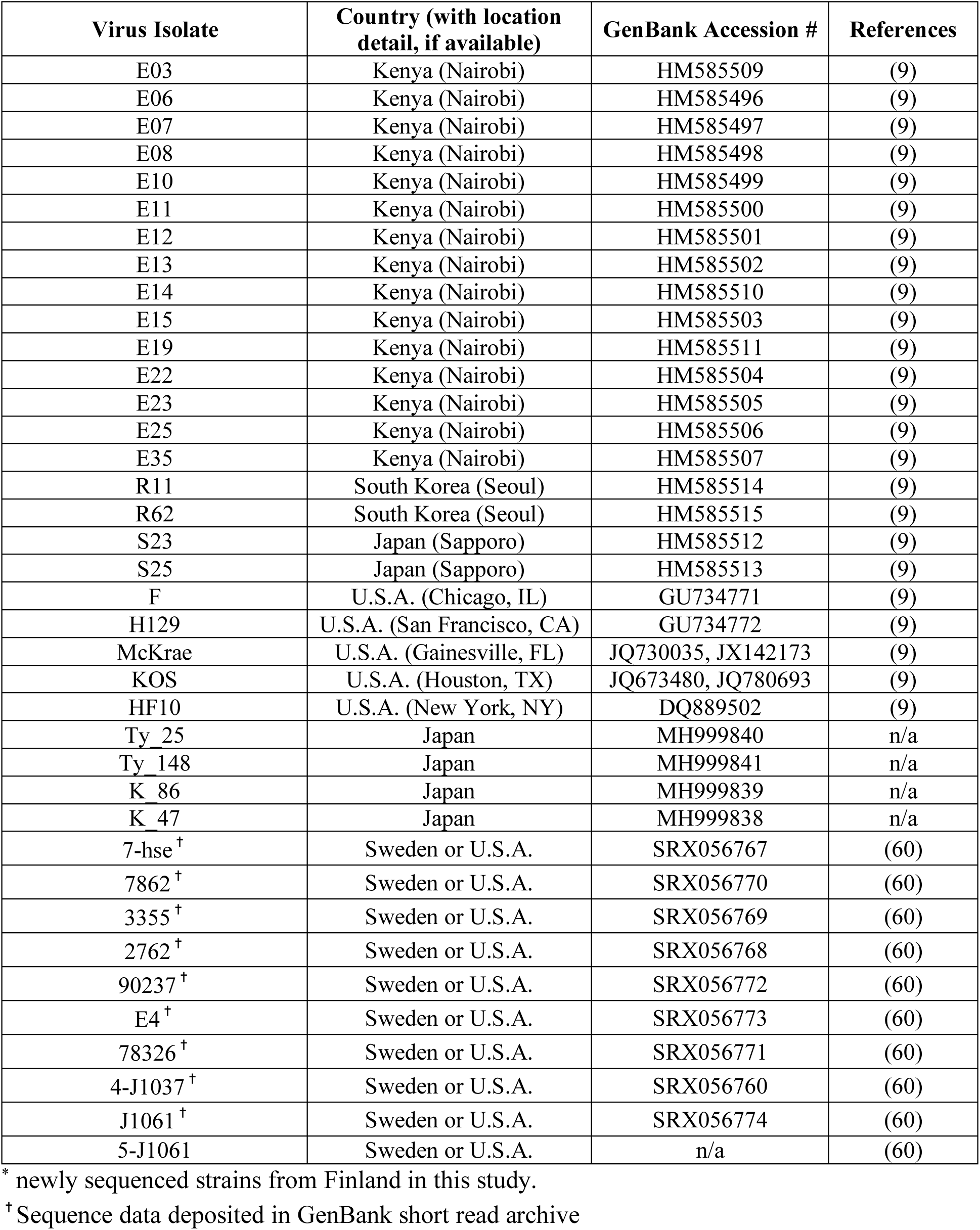
List of previously published HSV-1 genomes used for phylogenetic analyses

**Supplemental Table S2:** Number of nucleotide and amino acid (AA) differences observed in each set of Finnish or worldwide published HSV1 genomes.

| Gene* | ORF length in NT^ | protein length (#AA) | 10 Finnish # NT diff.^ | 10 Finnish # AA changes | 10 Finnish %AA diff. | 10 Finnish dN/dS | All HSV1 # NT diff. | All HSV1 # AA changes | All HSV1 %AA diff. | All HSV1 dN/dS ratio |
| --- | --- | --- | --- | --- | --- | --- | --- | --- | --- | --- |
| UL1 | 675 | 224 | 22 | 12 | 5.4 | 1.20 | 58 | 31 | 13.8 | 1.15 |
| UL2 | 1005 | 334 | 24 | 8 | 2.4 | 0.50 | 62 | 28 | 8.4 | 0.82 |
| UL3 | 675 | 224 | 13 | 5 | 2.2 | 0.63 | 43 | 19 | 8.5 | 0.79 |
| UL4 | 600 | 199 | 11 | 2 | 1.0 | 0.22 | 43 | 18 | 9.0 | 0.72 |
| UL5 | 2649 | 882 | 35 | 13 | 1.5 | 0.59 | 106 | 38 | 4.3 | 0.56 |
| UL6 | 2031 | 676 | 43 | 13 | 1.9 | 0.43 | 107 | 30 | 4.4 | 0.39 |
| UL7 | 891 | 296 | 24 | 6 | 2.0 | 0.33 | 47 | 17 | 5.7 | 0.57 |
| UL8 | 2253 | 750 | 44 | 18 | 2.4 | 0.69 | 124 | 62 | 8.3 | 1.00 |
| UL9 | 2556 | 851 | 21 | 5 | 0.6 | 0.31 | 110 | 32 | 3.8 | 0.41 |
| UL10 | 1422 | 473 | 25 | 5 | 1.1 | 0.25 | 81 | 30 | 6.3 | 0.59 |
| UL11 | 291 | 96 | 13 | 5 | 5.2 | 0.63 | 26 | 17 | 17.7 | 1.89 |
| UL12 | 1881 | 626 | 36 | 12 | 1.9 | 0.50 | 87 | 30 | 4.8 | 0.53 |
| UL13 | 1557 | 518 | 27 | 10 | 1.9 | 0.59 | 72 | 33 | 6.4 | 0.85 |
| UL14 | 660 | 219 | 10 | 6 | 2.7 | 1.50 | 27 | 15 | 6.8 | 1.25 |
| UL15 | 2208 | 735 | 22 | 3 | 0.4 | 0.16 | 80 | 16 | 2.2 | 0.25 |
| UL16 | 1122 | 373 | 13 | 3 | 0.8 | 0.30 | 55 | 19 | 5.1 | 0.53 |
| UL17 | 2112 | 703 | 30 | 12 | 1.7 | 0.67 | 108 | 39 | 5.5 | 0.57 |
| UL18 | 957 | 318 | 12 | 6 | 1.9 | 1.00 | 41 | 15 | 4.7 | 0.58 |
| UL19 | 4125 | 1374 | 39 | 7 | 0.5 | 0.22 | 139 | 40 | 2.9 | 0.40 |
| UL20 | 669 | 222 | 7 | 0 | 0.0 | 0.00 | 29 | 9 | 4.1 | 0.45 |
| UL21 | 1608 | 535 | 20 | 9 | 1.7 | 0.82 | 70 | 25 | 4.7 | 0.56 |
| UL22 | 2517 | 838 | 21 | 8 | 1.0 | 0.62 | 120 | 50 | 6.0 | 0.71 |
| UL23 | 1131 | 376 | 21 | 8 | 2.1 | 0.62 | 63 | 26 | 6.9 | 0.70 |
| UL24 | 810 | 269 | 18 | 7 | 2.6 | 0.64 | 60 | 28 | 10.4 | 0.88 |
| UL25 | 1743 | 580 | 19 | 4 | 0.7 | 0.27 | 74 | 25 | 4.3 | 0.51 |
| UL26 | 1908 | 635 | 40 | 14 | 2.2 | 0.54 | 92 | 34 | 5.4 | 0.59 |
| UL26.5 | 990 | 329 | 21 | 8 | 2.4 | 0.62 | 45 | 17 | 5.2 | 0.61 |
| UL27 | 2715 | 904 | 51 | 14 | 1.5 | 0.38 | 123 | 40 | 4.4 | 0.48 |
| UL28 | 2358 | 785 | 26 | 6 | 0.8 | 0.30 | 87 | 23 | 2.9 | 0.36 |
| UL29 | 3591 | 1196 | 54 | 10 | 0.8 | 0.23 | 154 | 33 | 2.8 | 0.27 |
| UL30 | 3708 | 1235 | 32 | 13 | 1.1 | 0.68 | 154 | 49 | 4.0 | 0.47 |
| UL31 | 921 | 306 | 15 | 5 | 1.6 | 0.50 | 34 | 13 | 4.2 | 0.62 |
| UL32 | 1791 | 596 | 20 | 2 | 0.3 | 0.11 | 85 | 28 | 4.7 | 0.49 |
| UL33 | 393 | 130 | 2 | 1 | 0.8 | 1.00 | 14 | 4 | 3.1 | 0.40 |
| UL34 | 828 | 275 | 15 | 6 | 2.2 | 0.67 | 37 | 14 | 5.1 | 0.61 |
| UL35 | 339 | 112 | 1 | 0 | 0.0 | 0.00 | 8 | 1 | 0.9 | 0.14 |
| UL36 | 9420 | 3139 | 166 | 66 | 2.1 | 0.66 | 1271 | 440 | 14.0 | 0.53 |
| UL37 | 3372 | 1123 | 41 | 13 | 1.2 | 0.46 | 140 | 60 | 5.3 | 0.75 |
| UL38 | 1398 | 465 | 18 | 8 | 1.7 | 0.80 | 70 | 30 | 6.5 | 0.75 |
| UL39 | 3414 | 1137 | 42 | 13 | 1.1 | 0.45 | 157 | 61 | 5.4 | 0.64 |
| UL40 | 1023 | 340 | 10 | 3 | 0.9 | 0.43 | 38 | 12 | 3.5 | 0.46 |
| UL41 | 1470 | 489 | 16 | 3 | 0.6 | 0.23 | 52 | 22 | 4.5 | 0.73 |
| UL42 | 1467 | 488 | 19 | 10 | 2.0 | 1.11 | 76 | 39 | 8.0 | 1.05 |
| UL43 | 1254 | 417 | 35 | 17 | 4.1 | 0.94 | 90 | 50 | 12.0 | 1.25 |
| UL44 | 1536 | 511 | 26 | 16 | 3.1 | 1.60 | 97 | 55 | 10.8 | 1.31 |
| UL45 | 519 | 172 | 4 | 3 | 1.7 | 3.00 | 22 | 8 | 4.7 | 0.57 |
| UL46 | 2157 | 718 | 33 | 15 | 2.1 | 0.83 | 111 | 65 | 9.1 | 1.41 |
| UL47 | 2082 | 693 | 29 | 6 | 0.9 | 0.26 | 81 | 29 | 4.2 | 0.56 |
| UL48 | 1473 | 490 | 18 | 4 | 0.8 | 0.29 | 69 | 25 | 5.1 | 0.57 |
| UL49 | 276 | 301 | 13 | 5 | 1.7 | 0.63 | 42 | 18 | 6.0 | 0.75 |
| UL49A | 906 | 91 | 2 | 0 | 0.0 | 0.00 | 8 | 3 | 3.3 | 0.60 |
| UL50 | 1116 | 371 | 23 | 9 | 2.4 | 0.64 | 65 | 26 | 7.0 | 0.67 |

| Gene* | ORF length in NT<sup>^</sup> | protein length (#AA) | 10 Finnish # NT diff.<sup>^</sup> | 10 Finnish # AA changes | 10 Finnish %AA diff. | 10 Finnish dN/dS | All HSV1 # NT diff. | All HSV1 # AA changes | All HSV1 %AA diff. | All HSV1 dN/dS ratio |
| --- | --- | --- | --- | --- | --- | --- | --- | --- | --- | --- |
| UL51 | 735 | 244 | 12 | 3 | 1.2 | 0.33 | 33 | 13 | 5.3 | 0.65 |
| UL52 | 3177 | 1058 | 43 | 9 | 0.9 | 0.26 | 129 | 41 | 3.9 | 0.47 |
| UL53 | 1017 | 338 | 16 | 4 | 1.2 | 0.33 | 36 | 12 | 3.6 | 0.50 |
| UL54 | 1539 | 512 | 11 | 4 | 0.8 | 0.57 | 73 | 34 | 6.6 | 0.87 |
| UL55 | 561 | 186 | 5 | 0 | 0.0 | 0.00 | 25 | 9 | 4.8 | 0.56 |
| UL56 | 705 | 234 | 8 | 4 | 1.7 | 1.00 | 34 | 19 | 8.1 | 1.27 |
| US1 | 1263 | 420 | 22 | 8 | 1.9 | 0.57 | 75 | 39 | 9.3 | 1.08 |
| US2 | 876 | 291 | 24 | 6 | 2.1 | 0.33 | 44 | 14 | 4.8 | 0.47 |
| US3 | 1446 | 481 | 14 | 5 | 1.0 | 0.56 | 57 | 21 | 4.4 | 0.58 |
| US4 | 717 | 238 | 26 | 13 | 5.5 | 1.00 | 68 | 36 | 15.1 | 1.13 |
| US5 | 279 | 92 | 10 | 7 | 7.6 | 2.33 | 25 | 17 | 18.5 | 2.13 |
| US6 | 1185 | 394 | 18 | 4 | 1.0 | 0.29 | 67 | 18 | 4.6 | 0.37 |
| US7 | 1173 | 390 | 28 | 12 | 3.1 | 0.75 | 119 | 57 | 14.6 | 0.92 |
| US8 | 1653 | 550 | 30 | 13 | 2.4 | 0.76 | 79 | 38 | 23.9 | 0.93 |
| US8A | 480 | 159 | 3 | 3 | 1.9 | 1.00 | 49 | 15 | 2.7 | 0.44 |
| US9 | 273 | 90 | 1 | 1 | 1.1 | 1.00 | 12 | 4 | 4.4 | 0.50 |
| US10 | 939 | 312 | 11 | 5 | 1.6 | 0.83 | 65 | 29 | 9.3 | 0.81 |
| US11 | 486 | 161 | 1 | 1 | 0.6 | 1.00 | 42 | 19 | 11.8 | 0.83 |
| US12 | 267 | 88 | 4 | 2 | 2.3 | 1.00 | 17 | 10 | 11.4 | 1.43 |
| RL1 | 747 | 248 | 25 | 11 | 4.4 | 0.79 | 422 | 90 | 36.3 | 0.27 |
| RL2 | 2328 | 775 | 44 | 20 | 2.6 | 0.83 | 180 | 91 | 11.7 | 1.02 |
| RS1 | 3897 | 1298 | 62 | 30 | 2.3 | 0.94 | 345 | 207 | 15.9 | 1.50 |

\* See **Supplemental Table S1** for list of strain names and genome accessions for comparisons of 10 Finnish and all published HSV1 genomes.
See **Supplemental Table S3** for list of strains excluded from selected gene alignments due to missing data in GenBank.
<sup>^</sup> Abbreviations: ORF, open reading frame; NT, nucleotide; AA, amino acid; diff., difference; %, percent; dN/dS, ratio of nonsynonymous to synonymous nucleotide changes

**Supplemental Table S3:** HSV-1 strains used to calculate nucleotide and amino acid (AA) differences in Table S2 and Figure 4.

| Gene | # of strains in nucleotide alignment | # of strains in AA alignment | Strains not used in nucleotide and AA alignments* |
| --- | --- | --- | --- |
| UL7 | 60 | 60 | B<sup>3</sup>x1.1, B<sup>3</sup>x1.2, B<sup>3</sup>x1.3, B<sup>3</sup>x1.4, B<sup>3</sup>x1.5 |
| UL12 | 61 | 61 | B<sup>3</sup>x1.3, B<sup>3</sup>x1.4, B<sup>3</sup>x1.5, RE |
| UL13 | 58 | 58 | E06, E25, E11, OD4, B<sup>3</sup>x1.3, B<sup>3</sup>x1.4, B<sup>3</sup>x1.5 |
| UL15 | 60 | 60 | B<sup>3</sup>x1.1, B<sup>3</sup>x1.2, B<sup>3</sup>x1.3, B<sup>3</sup>x1.4, B<sup>3</sup>x1.5 |
| UL19 | 61 | 61 | B<sup>3</sup>x1.3, B<sup>3</sup>x1.4, B<sup>3</sup>x1.5, RE |
| UL26.5 | 61 | 61 | B<sup>3</sup>x1.3, B<sup>3</sup>x1.4, B<sup>3</sup>x1.5, HF10 |
| UL41 | 61 | 61 | B<sup>3</sup>x1.3, B<sup>3</sup>x1.4, B<sup>3</sup>x1.5, OD4 |
| UL43 | 61 | 61 | B<sup>3</sup>x1.3, B<sup>3</sup>x1.4, B<sup>3</sup>x1.5, HF10 |
| UL48 | 60 | 60 | B<sup>3</sup>x1.1, B<sup>3</sup>x1.2, B<sup>3</sup>x1.3, B<sup>3</sup>x1.4, B<sup>3</sup>x1.5 |
| UL49A | 51 | 51 | B<sup>3</sup>x1.3, B<sup>3</sup>x1.4, B<sup>3</sup>x1.5, HF10, 66_2007, 369_2007, 160_1982, 1394_2005, 132_1998, 3083_2008, 2158_2007, 172_2010, 270_2007, 1319_2005 |
| UL55 | 59 | 59 | E35, E13, KUTy25, B<sup>3</sup>x1.3, B<sup>3</sup>x1.4, B<sup>3</sup>x1.5 |
| UL56 | 60 | 60 | B<sup>3</sup>x1.3, B<sup>3</sup>x1.4, B<sup>3</sup>x1.5, E35, HF10 |
| US1 | 60 | 60 | B<sup>3</sup>x1.1, B<sup>3</sup>x1.2, B<sup>3</sup>x1.3, B<sup>3</sup>x1.4, B<sup>3</sup>x1.5 |
| US8A | 61 | 61 | HF10, B<sup>3</sup>x1.3, B<sup>3</sup>x1.4, B<sup>3</sup>x1.5 |
| RL1 | 58 | 58 | B<sup>3</sup>x1.3, B<sup>3</sup>x1.4, B<sup>3</sup>x1.5, OD4, 1394_2005, CJ994, L2 |
| RL2 | 61 | 61 | B<sup>3</sup>x1.3, B<sup>3</sup>x1.4, B<sup>3</sup>x1.5, B<sup>3</sup>x1.2 |
| RS1 | 59 | 59 | B<sup>3</sup>x1.3, B<sup>3</sup>x1.4, B<sup>3</sup>x1.5, S23, CJ994, L2 |
| All other genes | 62 | 62 | B<sup>3</sup>x1.3, B<sup>3</sup>x1.4, B<sup>3</sup>x1.5 |

\* Strains were excluded due to missing data (e.g. due to sequencing gaps) or annotations in GenBank.

